# Association of RACK1 protein with ribosomes in *Plasmodium falciparum*

**DOI:** 10.1101/2021.09.21.461325

**Authors:** Jessey Erath, Sergej Djuranovic

## Abstract

The receptor for activated C-kinase 1 (RACK1), a highly conserved eukaryotic protein, is known to have many, varying biological roles and functions. Previous work has established RACK1 as a ribosomal protein, with defined regions important for binding ribosomes in both human and yeast cells. In *Plasmodium falciparum*, RACK1 has been shown to be required for parasite growth, however conflicting evidence has been presented about the RACK1 ribosome binding and its role in mRNA translation. Given the importance of RACK1 as a regulatory component of mRNA translation and ribosome quality control, the case could be made in parasites for either of the binary options: bound or unbound to the ribosome. Here we used bioinformatics and transcription analyses to describe *P. falciparum* RACK1 protein. Based on homology modeling and structural analyses, we generated a model of *P. falciparum* RACK1 protein. We created and explored mutant and chimeric human and *P. falciparum* RACK1 proteins binding properties to the human and *P. falciparum* ribosome. Wild type, chimeric and mutant RACK1 proteins suggest different binding characteristics for *P. falciparum* and human RACK1 proteins and different regions being involved in their ribosome association. The ribosomal binding of RACK1 variants in human and parasite cells shown here demonstrates that although RACK1 proteins have highly conserved sequences and structures across species, ribosomal binding is affected by species specific alterations to the protein. In conclusion, we show that in the case of *P. falciparum*, contrary to the structural data, RACK1 is found to be bound to ribosomes and in actively translating polysomes of *P. falciparum* cells.

## Background

RACK1 is a highly conserved, eukaryotic, seven-bladed WD β-propeller repeat scaffolding protein^1^. The WD domain proteins, fold into a β-propeller structure, is typically characterized by the repetition of glycine (G)–histidine (H) and tryptophan (W)-aspartic acid (D) dipeptide repeats^2^. They also happen to be incredibly abundant throughout the eukaryotic tree of life^3^. Proteins with WD domains are primarily associated with signalosome assembly^4^, providing the means for signal transduction, and there are as yet no WD domain proteins with intrinsic enzymatic capabilities^2^. Besides signal transduction, WD domains are involved in processes ranging from those involved in organism proliferation to virulence in lower eukaryotes^5^ or immune response in higher eukaryotes^1, 2,6^. RACK1 was indeed initially associated with protein kinase C signaling, the reason for its namesake^7^. However, RACK1 is now known to have many varying biological roles and functions, with the primary role of RACK1 as a ribosomal protein^8–10^. RACK1 has also been shown to be integral for multiple aspects of mRNA translation from efficient, cap-dependent mRNA translation initiation^8,9,11–14^ to recruitment of factors necessary for multiple mRNA/ribosome-associated quality control pathways^15–19^. The mammalian homolog having an *in vitro* half-life of 15 hours is stably associated with 40S ribosomal subunit and actively translating ribosomes^8^. This suggests a primary role of human RACK1 and its yeast ASC1 homolog as ribosomal proteins.

In *P. falciparum*, unlike yeast and mammalian cell lines^8^, RACK1 is required for parasite growth during IDC stages, with knockdown during ring stage result in growth arrest in the trophozoite stage^20^. Previous work has suggested that RACK1 is mainly _localized to parasitic cytoplasm during the schizont stage_ ^20^, but proteomic data has indicated that PfRACK1 might be exported to the erythrocyte cytoplasm for a yet to be define function^21^. What remains unclear are the functions of RACK1 in the parasite, given the lack of a PKC IIβ homolog^14^. Conflicting evidence has been provided regarding RACK1 ribosomal binding in *P. falciparum*. Structural data show that RACK1 does not co-purify with schizont 80S ribosomes^22,23^, while mass spectrometry of polysome profiling data paints a contradictory picture^24^. Authors of the initial structural study noted that loss of PfRACK1 binding to ribosomes may have been a result of different modes of ribosome binding or culturing conditions^22^. The authors of the second structural study suggested that the loss of RACK1 interaction may be driven by stage regulated binding, but not that of the EM sample preparation^23^. The mass spectrometry data^24^, however indicated a primarily ribosome-bound RACK1 and suggested binding of RACK1 to ribosome may be disrupted by experimental procedures. Lastly, the *in vitro* assessment of *P. falciparum* RACK1 ribosome binding was previously attempted with human ribosomes, however, authors were unable to exogenously produce and subsequently purify the parasite homolog due to suspected protein instability^8^. It is still unknown whether *P. falciparum* RACK1 protein binds to ribosomes and whether this ribosomal binding is conserved like in the case of mammalian and yeast homologs.

## Methods

### Transcriptional, bioinformatic and structural analysis of *P. falciparum* RACK1

In order to analyze the expression of *P. falciparum* RACK1 protein and potential interaction with ribosomes, we extracted transcriptional data for all ribosomal proteins from PlasmoDB^25^, excluding apicoplast and mitochondrial genes. Ribosomal proteins during the human stage were defined as those with RNAseq expression data in the minimum of the 80^th^ percentile during ring, early trophozoite, late trophozoite, schizont, gametocyte II, or gametocyte V^26^ or during sporozoite stage^27^. Ribosomal proteins expressed during the mosquito stage were defined as those with RNAseq expression data in the minimum of the 80^th^ percentile during ookinete^26^, oocyte, or sporozoite stage^27^.

Sequence analysis of RACK1 and ribosomal protein amino acid sequences was done by alignment using the Clustal Omega multiple sequence alignment tool^28^. The NCBI blastp suit was used to perform pair-wise sequence identity analysis on RACK1 amino acid sequences^29,30^. The secondary structure prediction was performed using the Quick2D tool in the MPI Bioinformatics Toolkit^31–37^. All structural alignments were performed using PyMOL (Version 2.0 Schrödinger, LLC.)^38^. The resulting *de novo* PfRACK1 model was added to the previously generated *P. falciparum* 80S structure (3JBO)^23^. First, the *de novo* PfRACK1 model was structurally aligned to the human RACK1 protein found in the previously generated *H. sapiens* 80S structure (3JAG)^39^. The *P. falciparum* 80S structure was then aligned to the *H. sapiens* 80S structure. Interacting residues between the 80S ribosome and RACK1 protein were those found to be within a 3-5 Å range. Electrostatic maps were generated using PyMOL APBS Electrostatics plugin^40^.

### De novo modeling of P. falciparum RACK1

The RACK1 variants were modeled using SWISS-MODEL^41–45^. Briefly, the RACK1 variant amino acid sequences were entered into the SWISS-MODEL server. The SWISS-MODEL software generates a list of potential structural templates based on the input. The software then aligns and models the input RACK1 variants on the selected templates and subsequently evaluates the quality of the model. Templates are then selected initially based on the GMQE score, coverage, and resolution. A final template is then selected based on the QMEAN score. Here, all RACK1 variants aligned and modeled well with the 4aow.1.A template, X-ray crystallography generated structure of human RACK1^28^.

### RACK1 variant construction

The PfRACK1 gene was codon-optimized for expression in human cells. The HsRACK1 gene was cloned from cDNA. None of the expressed RACK1 variants have the typical intron that appears to be conserved in the gene. Based on bioinformatics and structural analysis, RACK1 variants were designed. The swapping of the N-terminal regions and other mutants were generated by mutagenesis PCR, whereby the 5’ and 3’ PCR products were produced overlapping in the mutated regions. These products were combined by stitching PCR and cloned into the desired expression vectors. Clones of vectors with RACK1 variants were confirmed by Sanger sequencing.

### Mammalian cell culture, lentivirus production, and transduction

Human embryonic kidney (HEK) 293T cells were cultured in Dulbecco’s modified Eagle’s medium (DMEM) (Gibco) and supplemented with 10% fetal bovine serum, 5% penicillin and streptomycin (Gibco), and L-glutamine (Gibco). HAP1 cells were cultured in Iscove’s Modified Dulbecco’s Medium (IMDM) (Gibco) and supplemented with 10% Fetalgro® Bovine Growth Serum (RMBIO), 5% penicillin and streptomycin (Gibco), and L-glutamine (Gibco). Cells were grown at 37 °C with 5% CO_2_.

The You Lab (Washington University, St. Louis, MO) generously provided tools for lentiviral production. The RACK1 variants to be expressed were cloned into the pCDH vector using the XbaI and NotI restriction sites and confirmed by Sanger sequencing. Plasmid DNA from pCDH-RACK1, pCMV-VSVG vector (addgene 8454), and psPAX2 vector (addgene 12260) was isolated from bacterial midipreps (Invitrogen). Before transfection (20-24 hours), 1.5 million HEK 293T cells in a 6 cm dish. The packaging vectors, pCMV-VSVG and psPAX2, were combined in a ratio of 1:9. For transfection, 3 μg of the expression vector and 3 μg of combined packaging vectors were also combined in 400 μL of OPTI-MEM (Gibco). To this, 18 μL (1:3 ratio of DNA to transfection reagent) of X-tremeGENE™ 9 DNA transfection reagent (MilliPore Sigma). The mixture was incubated for 30 mins at room temperature. The medium of HEK 293T cells was replaced with pre-warmed regular growth medium. The transfection mixture was added, and cell put back in the incubator at 37 °C with 5% CO_2_. On the next day, 1 mL of fetal bovine serum (FBS) was added. The medium containing virus was collected at 48 hours and 72 hours after transfection. The supernatant containing the virus was centrifugated at 15,000 x g and then filtered using 0.45 μm filter to remove any 293T cells or debris. The virus was stored at 4 °C for up to 1 week or kept at -80 °C for long term storage, and freeze-thawing was avoided.

For lentiviral transduction, HAP1 cells were seed such that they will be about 60-70% confluent at the time of transduction. To assess viral titer, cells were plated in 12-well plates. However, a 1:4 dilution was found to be sufficient enough for an almost 100% transduction. At 20-24 hours after plating, polybrene was added to a final concentration of 8μg/mL to cells and the desired amount of lentivirus was added. At 48 hours after transduction, cells were re-plated and selected with 3 μg/mL puromycin for 24 hours, two to three times.

### Parasite cell culture and transfection

*P. falciparum* Dd2 was cultured at 2-5% hematocrit in O^+^ erythrocytes in Malaria Culture Medium (MCM): RPMI 1640 supplemented with 5 g/L Albumax II (Gibco), 0.12 mM hypoxanthine (1.2 mL 0.1M hypoxanthine in 1 M NaOH), 10 μg/mL gentamicin^46^. Cultures were grown statically in candle jar atmosphere. As required, cultures were synchronized with 5% (wt/vol) sorbitol. Asynchronous *P. falciparum* Dd2 parasites were washed twice in 15 mL incomplete Cytomix (25 mM HEPES, 0.15 mM CaCl_2_, 5 mM MgCl_2_, 2 mM EGTA, 120 mM KCl, 5 mM K_2_HPO_4_, 5 mM KH_2_PO_4_) and resuspended in a total 525 μL with 100 μg of maxi-prep plasmid DNA dissolved in incomplete cytomix (125 μL packed iRBCs and 400 μL DNA/incomplete cytomix), The parasites were transfected in Bio-Rad Gene Pulser cuvette (0.2 cm), 0.31 kV, 950 up, infinity resistance. Selection (10 nM WR99210 or 2 μM DSM-1) was added to the parasite 48 h after transfection and used to select resistant parasites^47^.

### Saponin lysis of infected red blood cells (iRBCs)

The cell iRBCs were resuspended in two volumes of PBS containing 0.15% saponin, and incubated on ice for 10 min, with vigorous mixing every 3 min. Afterward, the samples were centrifuged 7000g, 5 min, 4°C, and the pellets were washed three times more with the same buffer.

### Parasite staining and confocal microscopy with airyscan

To examine localization of RACK1 variants in the parasite, confocal microscopy using the Zeiss LSM 880 Confocal with Airyscan was performed. Primarily late trophozoite, early schizont stage parasites were treated for 6 hours with E64 to inhibit parasite release^48,49^. Parasites were then washed twice using a parasite imaging medium (RPMI without phenol red and with hypoxanthine) containing E64. Parasites were incubated for 30 min with Hoechst stain to label DNA and BODIPY™ TR ceramide to label lipids. Parasites were again washed 3 times with an imaging medium containing E64. Cultures were dotted onto slides and sealed beneath coverslips. Brightfield images and z-stacks were collected using a Zeiss LSM 880 Confocal with Airyscan confocal microscope using a 40X oil objective. DNA was visualized by Hoechst stain (diode: 405 nm, ex: 350 nm, em: 461 nm), RACK by mNeonGreen (laser: 488 nm, ex: 506, em: 517), and lipid bilayers by BODIPY™ TR ceramide (laser: 561 nm, ex:589/em:617) Image analysis and generation was done using Imaris (Oxford Instruments) and Fiji ImageJ software.

### Crude ribosome preparation and polysome profiling

Crude ribosome pellets and polysome profiling in *P. falciparum* cells was done based on previous work^24^. Parasites were grown to 7-10% parasitemia in 1.5 to 2.0 mL of packed iRBCs. Parasites were treated with culture medium containing 200 μM cyclohexamide (CHX) for 10 minutes at 37°C. They were then washed three times with 1X PBS + 200 μM CHX. Samples were then flash-frozen in liquid nitrogen and stored at -80°C overnight. To the frozen samples, 2 volumes of lysis buffer (25 mM potassium HEPES pH 7.2, 400 mM potassium acetate, 15 mM magnesium acetate, 1% IGEPAL® CA-360, 200 μM CHX, 1 mM DTT, 1 mM AEBSF, 40 U/mL RNase inhibitor) and 0.5 mg of acid washed glass beads (Sigma #G8772) were added. Samples were rotated end-over-end at 4°C for 10 minutes. Samples were centrifugated for 5 minutes at 1000xg. Supernatants were transferred to a new tube and centrifugated at 14000 x g for 10 minutes. The supernatant was layered over 1 M sucrose cushion (1 M sucrose, 25 mM potassium HEPES pH 7.2, 400 mM potassium acetate, 15 mM magnesium acetate, 200 μM CHX, 1 mM DTT, 1 mM AEBSF, 40 U/mL RNase inhibitor) in a ratio of 2 mL lysate to 1 mL sucrose cushion. The balanced tubes were then placed in a TLA 100.3 rotor and centrifuged for 30 minutes at 335,000 x g at 4°C using an Optima™□ Max-XP Beckman Coulter ultracentrifuge. The supernatants were removed. The crude ribosome pellets were washed with 500 μL wash buffer (25 mM potassium HEPES pH 7.2, 400 mM potassium acetate, 15 mM magnesium acetate, 200 μM CHX, 1 mM DTT, 0.1 mM AEBSF, 10 U/mL RNase inhibitor) three times. After washing, the crude ribosome pellets were then resuspended in 200 μL of lysis buffer. The resuspensions were again layered over 1 mL of 1 M sucrose cushion and the pelleting repeated as above. Washing was repeated as previous. The crude ribosome pellets used in polysome profiling were then resuspended in 200 μL of ribosome resuspension buffer and, if not immediately used, flash frozen in liquid nitrogen and store at -80°C for later use. Crude ribosome pellets used for immunoblotting were resuspended in sample buffer at 37°C for 20 minutes, boiled at 95°C for 5 minutes, and chilled on ice. Sucrose gradients were generated using polysome profiling buffer (25 mM potassium HEPES pH 7.2, 400 mM potassium acetate, 15 mM magnesium acetate, 200 μM CHX, 1 mM DTT, 1 mM AEBSF) with 10%, 20%, 30%, 40%, and 50% sucrose layers in 14 mm x 89 mm polyallomer centrifuge tubes (Beckman #331372). Total RNA of resuspended crude ribosomes was measured to ensure a minimum of 300 μg total was loaded onto the gradient. Samples were layered over the gradient and balanced using polysome profiling buffer. The gradients were centrifuged at 35,000 RPM (acceleration 3, deceleration 4) for 2 hours and 40 minutes at 4°C using an Optima™□ L-100 XP Beckman Coulter ultracentrifuge in a SW41 swing bucket rotor. Centrifuged gradients were then fractionated using a density gradient fractionation system (BRANDEL #BR188176). Samples were collected into 1.7 mL microcentrifuge tubes with an equal volume of Ribozol™ (VWR #VWRVN580) with 1% SDS for RNA or 3 volumes of 20% TCA in MilliQ water for protein. Samples were mixed by vortexing. RNA was stored at - 80°C if not isolated immediately. RNA was isolated following manufacturer protocol and used in RNA gel electrophoresis. Protein samples were stored or incubated at 4°C overnight to precipitate protein. Samples were then centrifuge at 20,000 x g, 4°C for 30 minutes. Supernatant was removed. The pellets were washed with 100% acetone twice. Pellets were then solubilized in sample buffer for western blot analysis.

Crude ribosome pellets and polysome profiling in mammalian cells was done based on the previous work^50^. HAP1 cells were grown to 80-90% confluency in a 10 cm dish. Cells were washed three times with 5 mL of ice-cold PBS + 200 μM CHX. A 400 μL volume of lysis buffer (20 mM Tris-HCl pH 7.4, 150 mM NaCl, 5 mM MgCl_2_, 1% v/v Triton X-100, 200 μM CHX, 1 mM DTT, 1 mM AEBSF, 40 U/mL RNase inhibitor) was then added dropwise to the place and cells were scraped off and collected into a 1.7 mL microcentrifuge tube. The lysate was pipetted several times and incubated on ice for 10 minutes. The samples were then centrifugated at 20,000 x g for 10 minutes at 4°C to remove cell debris. The lysate was transferred into a new tube. If not used immediately, lysates were flash-frozen in liquid nitrogen and stored at -80°C. Crude ribosome pellets were generated by layering 200 μL of lysate over 0.9 mL of 1M sucrose cushion (1 M sucrose, 20 mM Tris-HCl pH 7.4, 150 mM NaCl, 5 mM MgCl_2_, 200 μM CHX, 1 mM DTT, 1 mM AEBSF, 40 U/mL RNase inhibitor). The balanced tubes were then placed in a TLA 100.3 rotor and centrifuged for 1 hour at 100,000 x g at 4°C using an Optima™□ Max-XP Beckman Coulter ultracentrifuge. The supernatants were removed. The crude ribosome pellets were washed with 500 μL wash buffer (20 mM Tris-HCl pH 7.4, 150 mM NaCl, 5 mM MgCl_2_, 0.1% v/v Tween 20, 200 μM CHX, 1 mM DTT, 0.1 mM AEBSF, 10 U/mL RNase inhibitor) three times. Crude ribosome pellets used for immunoblotting were resuspended in sample buffer at 37°C for 20 minutes, boiled at 95°C for 5 minutes, and chilled on ice. Polysome profiling was performed by layering clarified HAP1 cell lysates over sucrose gradient generated with polysome profiling buffer (20 mM Tris-HCl pH 7.4, 150 mM NaCl, 5 mM MgCl_2_, 200 μM CHX, 1 mM DTT, 1 mM AEBSF, 40 U/mL RNase inhibitor) and centrifugation performed as above. Samples were collected and isolated at above for RNA gel electrophoresis and western blot analysis.

### HA-immunoprecipitation and immunoblotting

Mixed *Plasmodium falciparum* Dd2 parasites were harvested at 7-10% parasitemia. For examination of RACK1 variant expression, parasites were released from RBCs by saponin lysis as previously described. Parasites were then lysed using passive lysis buffer (Promega #E1910) and the total protein concentration was calculated by RC/DC kit (BioRad #5000122). For each reaction, 25 μL of magnetic HA beads was washed twice in 1X PBS with 0.1% Tween 20 and then twice in binding buffer (50 mM Tris pH7.5, 150 mM NaCl, 1% IGEPAL CA-630, 5% glycerol, and protease inhibitors (Thermo Fisher #A32955): apoptin, leupeptin, bestatin, E-64, AEBSF, pepstatin A). A total of 500 μg of protein was loaded onto the HA beads and bound overnight at 4°C. The beads were then washed three times with 1 mL of wash buffer (50 mM Tris pH7.5, 150 mM NaCl, 1% IGEPAL CA-630, 5% glycerol). Beads were then eluted in 50 μL of sample buffer and examined by western blot analysis. Samples were loaded onto SDS-PAGE gels, running at 15 W, 1h 10 mins. The gels were then transferred to PVDF membrane using semi-dry transfer method running at 25 V for 40 minutes. The PVDF membranes were blocked in 5% milk in PBS + 0.1% Tween 20 (PBST) for 1 hour at room temperature or overnight at 4°C. The membranes were probed with diluted primary antibody (see **SUPPLEMENTARY TABLE 4.3** for antibodies) in PBST with 5% milk. After incubation with the primary antibody, the PVDF membranes were washed three times for 5 min in PBST. Membranes incubated with HRP-conjugated primaries were incubated and washed similarly, washing with PBST, and then proceeding to imaging instead of secondary. The membranes were then incubated with HRP-conjugated secondary antibody diluted 1:2500 in PBST with 5% milk for 1 hour at room temperature. The membranes were washed as above. Then rinsed with PBS. Prepare Working Solution by mixing equal parts of the Stable Peroxide Solution and the Luminol/Enhancer Solution (34577 SUPERSIGNAL WEST PICO PLUS, 34096 SUPERSIGNAL WEST FEMTO MAXIMUM SENSITIVITY SUBSTRATE, respectively). We incubated the blot in Working Solution for 5 minutes. Remove the blot from Working Solution and drain excess reagent. Afterward, we took images were generated by BioRad Molecular Imager CHemiDoc XRS System with Image Lab software.

### qPCR Analysis of polysome rRNA and rRNA bound to RACK1 variant

RNA isolated from polysome profiling fractions were analyzed by qPCR. Using the NanoDrop 2000c, 50 ng of total RNA was used to generate cDNA via the iScript Advance cDNA Kit for RT-qPCR (BioRAD #1725037), for each sample including no reverse transcriptase and no template controls following the manufacturer’s protocol. The iTaq Universal SYBR Green Supermix (BioRAD #1725121) protocol was used for qRT-PCR on the CFX96 Real-Time system with Bio-Rad CFX Manager 3.0 software^51^. The 2^-Ct^ value were calculated and plotted for each fraction.

HA-immunoprecipitation was performed on parasite crude ribosome pellets, both previously described. The total protein concentration in each crude ribosome pellet was calculated by RC/DC kit (BioRad #5000122). The beads were then blocked with polysome parasite lysis buffer containing 4% BSA and 0.5 μg/mL *S. cerevisiae* tRNAs for one hour at 4°C while rotating. A total of 680 μg of protein was loaded onto the HA beads and bound for one hour at 4°C while rotating. The beads were then washed three times with 1 mL of polysome wash buffer. Beads were split in half. For protein, beads were then eluted in 35 μL of sample buffer and examined by western blot analysis as previously described. For RNA, beads were eluted with 400 μL Trizol + 1% SDS at 36°C while shaking. RNA was isolated by per the manufacturer’s protocol. Using the NanoDrop 2000c, 50 ng of total RNA was used to generate cDNA via the iScript Advance cDNA Kit for RT-qPCR (BioRAD #1725037), for each sample including no reverse transcriptase and no template controls following the manufacturer’s protocol. The iTaq Universal SYBR Green Supermix (BioRAD #1725121) protocol was used for qRT-PCR on the CFX96 Real-Time system with Bio-Rad CFX Manager 3.0 software^51^. The ΔCt was calculated by subtracting the Dd2 parent line values from each sample. The ΔΔCt value was then calculated by normalization to the wild-type PfRACK1 variant. Plots were then generated using the calculated 2^-ΔΔCt^ values for each variant.

## Results

### Sequence analysis of RACK1 proteins

As previously mentioned, RACK1 is a highly conserved, eukaryotic scaffolding protein with a seven-bladed WD β-propeller repeat structure^1^. We performed bioinformatic analysis on selected set of organisms including other single-celled parasitic organisms, a human malaria vector, the model organisms *Drosophila melanogaster, Saccharomyces cerevisiae*, and the human host (**FIGURE 1** and **TABLE 1**). Expectedly, the *Plasmodium spp.* share the highest sequence identity with *Plasmodium falciparum* Dd2 strain RACK1 (PfRACK1) protein, whereby human-infective strains *P. vivax* P01 and *P. knowlesi* strain H show only minor differences. This is followed by the murine parasite species *P. chabaudi chabaudi*, *P. yoelii yoelii* 17X, and *P. berghei* ANKA. The RACK1 homolog expressed by *Toxoplasma gondii*, a fellow Apicomplexan whose genes and functions are often compared with *Plasmodium spp.*, is also quite similar at 67.30% (**TABLE 1**). Interestingly, the *H. sapiens* RACK (HsRACK1) homolog has a shared identity of almost 60%, higher than other single-celled parasites (*T. brucei brucei* at 51.30% and *L. brazilliensis* at 41.97%) and *S. cerevisiae* (ScRACK1, 42.99%) that one might anticipate sharing greater sequence identity. Lastly, the model organism *D. melanogaster* and malaria vector *A. gambiae* also share relatively high sequence identity at 58.15% and 56.45%, respectively.

**FIGURE 1.**
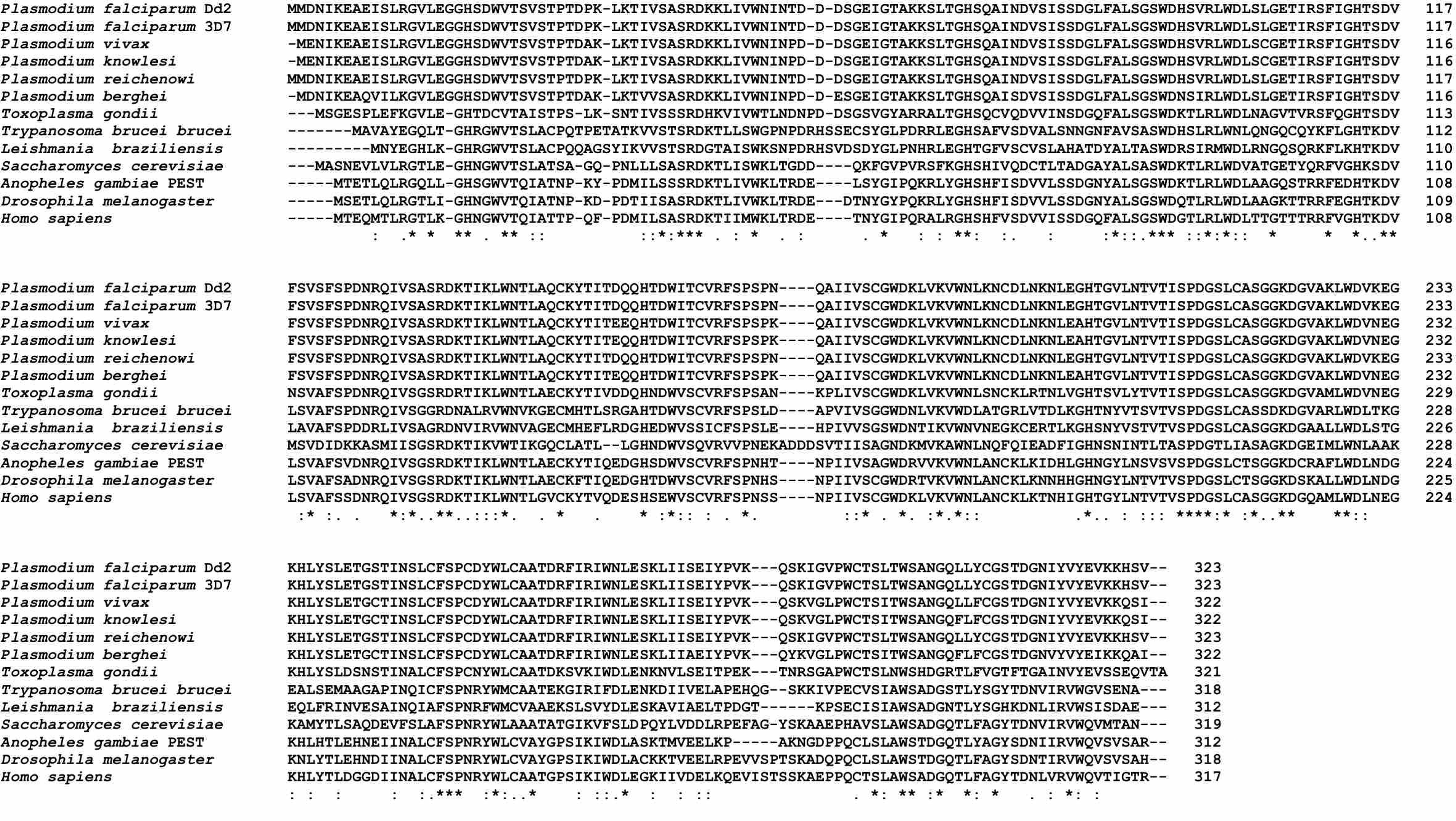
CLUSTAL OMEGA ALIGNMENT OF RACK1 HOMOLOGUE PROTEIN SEQUENCES FROM SELECTED ORGANISMS. The *Plasmodium falciparum* RACK1 protein sequence was aligned to selected organisms to compare with closely related, human-infective species (*P. vivax*, *P. knowlesi*, *P. reichenowi*) as well as murine model species (*P. berghei*). We also sought to examine conservation with the often compare parasite *T. gondii* and other eukaryotic parasites (*T. brucei brucei*, *L. braziliensis*). Two model organisms, one unicellular (*S. cerevisiae*) and the other multicellular (*D. melanogaster*), were also included. Lastly, the human host (*H. sapiens*) and vector (*A. gambiae* PEST) were also examined, with the human host being a focal point of the analysis. Clustal Omega consensus symbols: Asterisks (*) indicates fully conserved resdue. Colon (:) indicates conservation between residues of strongly similar properties (approximation of > 0.5 in the Gonnet PAM 250 matrix). Period (.) indicates conservation between residues of weakly similar properties (approximation of =< 0.5 and > 0 in the Gonnet PAM 250 matrix).

**TABLE 1.** ANALYSIS OF RACK1 SEQUENCE IDENTITIES VERSUS *P. FALCIPARUM* DD2 STRAIN. Sequence identities were determined using protein basic local alignment search tool to align protein sequences with P. falciparum Dd2 RACK1 protein sequence for each organism.

| **Organism** | **Identity (%)** |
| --- | --- |
| *P. falciparum* 3D7 | 100.00 |
| *P. reichenowi* CDC | 100.00 |
| *P. vivax* P01 | 95.03 |
| *P. knowlesi* strain H | 95.03 |
| *P. chabaudi chabaudi* | 91.93 |
| *P. yoelii yoelii* 17X | 91.61 |
| *P. berghei* ANKA | 91.30 |
| *T. gondii* | 67.30 |
| *H. sapiens* | 59.55 |
| *D. melanogaster* | 58.15 |
| *A. gambiae* PEST | 56.45 |
| *T. brucei brucei* TREU927 | 51.30 |
| *S. cerevisiae* | 42.99 |
| *L. braziliensis* MHOM/BR/75/M2904 | 41.97 |

Bioinformatic analysis of HsRACK1 and ScAsc1 proteins show that they share a 53.82% sequence identity, with HsRACK1 and PfRACK1 having higher sequence identity (59.55%). A multiple sequence alignment combined with secondary structure analysis of these RACK1 proteins was performed to further determine what features, if any, would affect the binding of PfRACK1 to the 40S ribosome subunit. The analysis demonstrates a conservation of the β-sheet blades and loop features and positioning (**FIGURE 2A**). Previously described residues, such as the RDK motif vital to ribosome binding (**FIGURE 2A** **MUTANT 3**), are conserved^1,9,10,52^. However, there are multiple regions within the N-terminus whereby significant residue changes result in charges differences that could affect RACK1 ribosomal binding. Some of these changes have been subtle (**FIGURE 2A** **MUTANT 1**), however others are more drastic, such as changes in or introductions of charge (**FIGURE 2A** **MUTANT 2 AND 5**) or loop length and residue polarity (**FIGURE 2A** **MUTANT 4**). The C-terminal of the proteins appears to be significantly more conserved except so-called “knob” region located in the sixth β-sheet blade suggested to be a species-specific region offering differences in translational control^53^. This region was shown to be dispensable in yeast^52^. This sequences is FAGYS in yeast, STSS in humans, and HQS in *P. falciparum*. Regardless, previous work suggests that the loop region has evolved to accommodate differences in eIF6 binding and 5’ polyA leader sequence usage by different eukaryotic organisms, thereby not necessarily being important of ribosome binding^53^.

**FIGURE 2.**
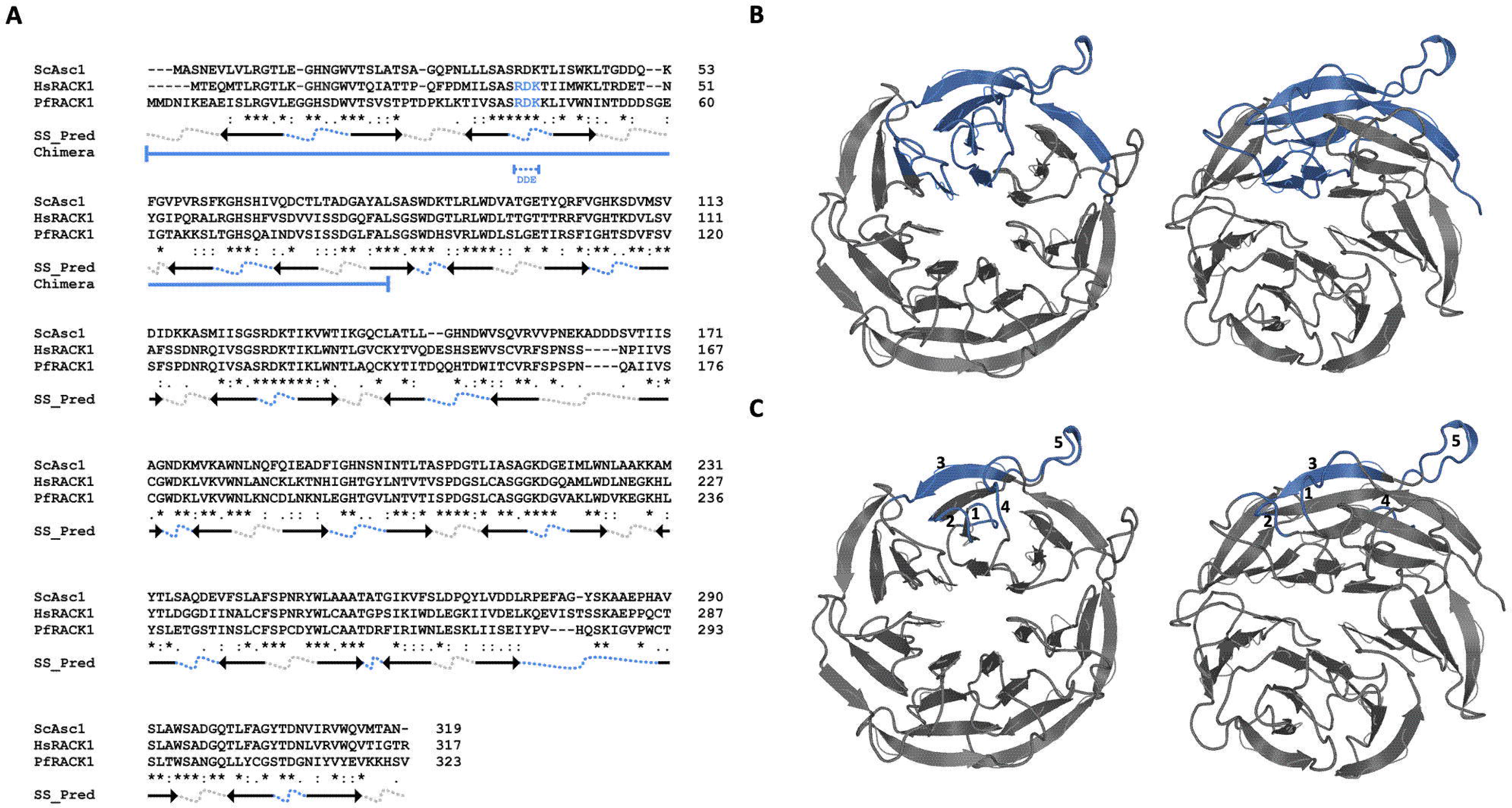
COMPARATIVE BIOINFORMATIC ANALYSIS OF RACK1 HOMOLOGS IN *S. CEREVISIAE* (SCASC1), *H. SAPIENS* (HSRACK1), AND *P. FALCIPARUM* (PFRACK1) HOMOLOGS. (A) Sequence alignment of yeast Asc1, human RACK1, and *P. falciparum* RACK1 amino acid sequences generated by Clustal Omega. The SS_Pred is the secondary structure prediction using MPI Bioinformatics toolkit Quick2D tool. Arrows: beta strands. Arrows represent beta strands, with heads pointing from the N-terminal to C-terminal direction showing the orientation of the beta strand in the β-propeller. Dotted lines represent loops found between beta strands where grey notates solvent-facing loops and blue indicates ribosome-facing loops. Chimera (χ): proteins generated by region exchanged between human and parasite RACK1 proteins shown in solid blue lines. RDK→DDE mutant indicated in blue text and blue dashed line. Clustal Omega consensus symbols: Asterisks (*) indicates fully conserved resdue. Colon (:) indicates conservation between residues of strongly similar properties (approximation of > 0.5 in the Gonnet PAM 250 matrix). Period (.) indicates conservation between residues of weakly similar properties (approximation of =< 0.5 and > 0 in the Gonnet PAM 250 matrix). **(B-C)** A SWISS MODEL generated de novo model of PfRACK1 displaying **(B)** chimeric region and **(C)** mutated regions in blue. Left: Ribosome facing surface. Right: 25-degree rotation.

### Modeling of *P. falciparum* RACK1 protein

To get further insights in differences and similarities between PfRACK1 and HsRACK1 proteins we created a model of PfRACK1 protein (**FIGURE 3**). The model of PfRACK1 protein was generated based on the sequence alignments (**FIGURE 1** **and** **FIGURE 2**) and homology modeling using SWISS-MODEL^41–45^. HsRACK1 has been previously shown to bind to the human 40S ribosome, with emphasis placed on helices 39 and 40 of the 18s rRNA and the N-terminal region of RACK1^54^. We checked whether this feature is preserved in *P. falciparum* RACK1 protein and displayed PfRACK1 as previously published for HsRACK1 interaction with the human 80S ribosome^54^ (**FIGURE 3A**). The helices 39 and 40 of the parasite 18S rRNA reveal the potential for highly similar interactions between the PfRACK1 protein and the *P. falciparum* 18S rRNA. Previous work also examined ribosomal proteins in this region: RPS16 (uS9), RPS17 (eS17), and RPS3 (uS3) (**FIGURE 3B–C**), which are also in close proximity. Therefore, residues that may interact with these proteins must also be taken into consideration. Those residues on PfRPS16 and PfRPS17 within 3-5 Å range are also highly conserved (**SUPPLEMENTAL FIGURE 1**), and therefore are not expected to hinder binding. The residues of PfRPS3 are also highly conserved, however, it appears that there may be a C-terminal truncation of 18 residues when compared with HsRPS3 that may impact interaction of the disordered region of the protein (**SUPPLEMENTAL FIGURE 1)**. Overall, the PfRACK1 model data shows high sequence and structural conservation of the PfRACK1 protein compared with HsRACK1 and the possibility of PfRACK1 to bind to the ribosome.

**FIGURE 3.**
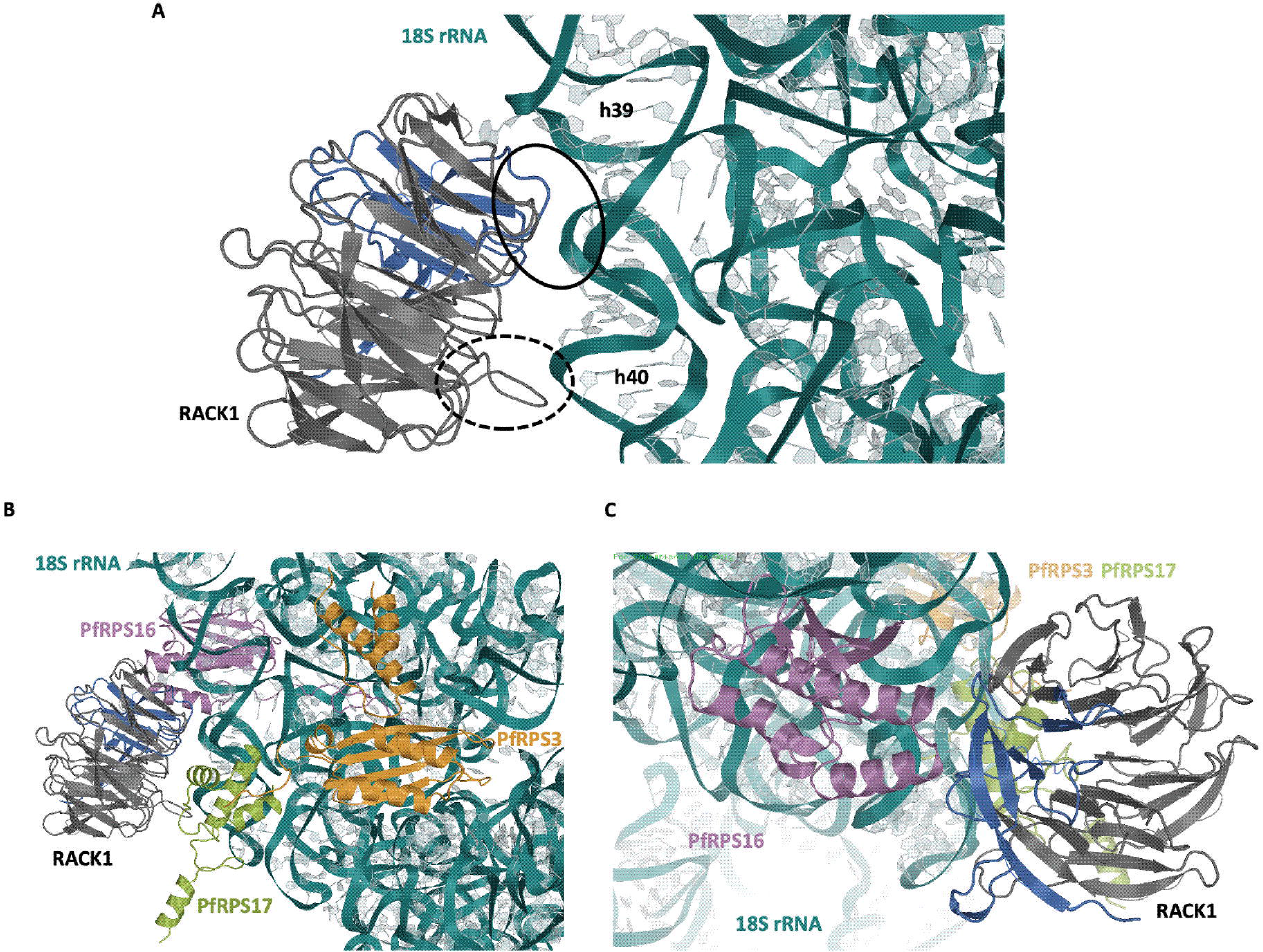
STRUCTURAL ANALYSIS OF THE PFRACK1 PROTEIN BINDING TO *P. FALCIPARUM* 40S RIBOSOME SUBUNIT. All structures displayed are P falciparum proteins/RNA. **(A)** The structure of the *P. falciparum* 80S ribosome (RCSB PDB ID: 3jbo) was structurally aligned with the human 80S ribosome (RCSB PDB ID: 3jag). The de novo model of PfRACK1 generated by SWISS-MODEL was structurally aligned with that of the human 80S bound RACK1. The view shown is that which been previously published arguing for the importance of the regions noted in RACK1 binding to the ribosome. Solid circle: region vital for HsRACK1:Hs80S binding. Dotted circle: Loop region that varies highly species to species, and while not required, may influence binding. **(B)** The same view as above, however ribosomal binding proteins with potential interactions are shown. **(C)** A 180-degree flipped view to show additional RPS16 (uS9) and RACK1 interactions. Note: Region in blue is that when exchanged between human and P. falciparum enabled binding of parasite RACK1 to the human 40S ribosome.

#### PfRACK1 expression follows other ribosomal proteins

We performed a transcriptional analysis of PfRACK1 to compare its expression profile over the different parasite life stages and correlation with expression of *P. falciparum* ribosomal proteins. RNAseq data for all available *P. falciparum* proteins for the ring, early trophozoite, late trophozoite, schizont, gametocyte II, gametocyte V, ookinete, oocyst, and sporozoite stages from the previous study^26^ was collected from PlasmoDB^25,26^. A violin plot of all data was generated with the individual data points for the ribosomal protein plotted. The RNAseq data points for RACK1 were then plotted in red to compare the RNA expression of RACK1 with the ribosomal proteins vs. all parasite proteins. Our analysis shows that the PfRACK1 gene, like other ribosomal proteins, is highly expressed compared to the average total protein transcript expression at each stage and in a pattern similar to other ribosomal proteins (**FIGURE 4**). Interestingly, PfRACK1 expression drops by approximately one log_10_ from the early trophozoite stage where translation is high to when the parasites transition into the schizont stage. Thus, the data suggest that PfRACK expression follows that of other ribosomal proteins and significant changes in expression are seen during stages where protein synthesis dramatically increases or decreases.

**FIGURE 4.**
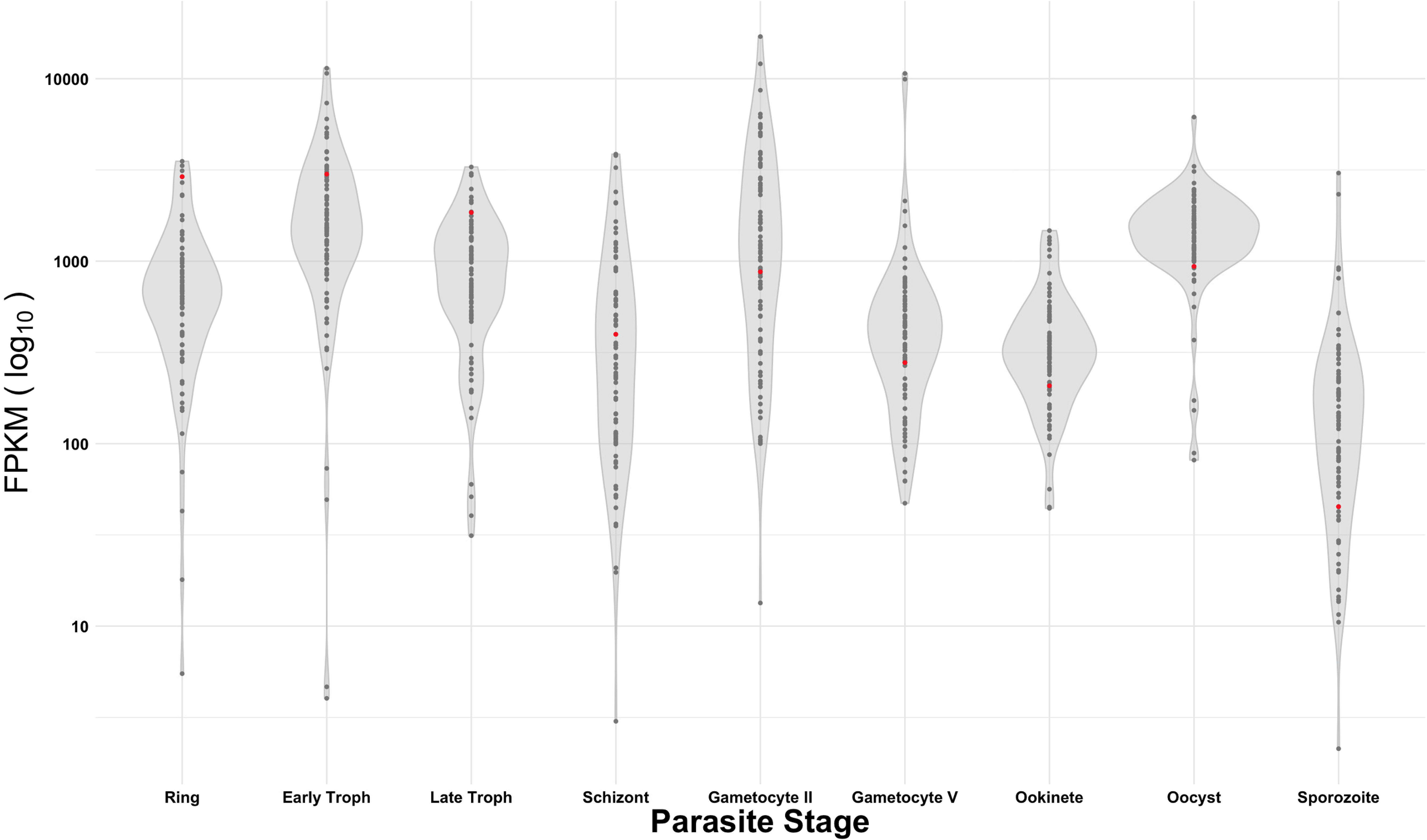
TRANSCRIPTIONAL ANALYSIS OF PFRACK1 IN COMPARISON WITH OTHER PARASITE PROTEINS THROUGHOUT THE *P. FALCIPARUM* LIFE CYCLE. RNASeq data from PlasmoDB showing all proteins (violin plot), ribosomal protein (dot plot), and PfRACK1 (red dot) transcript expression in *P. falciparum* 3D7 throughout the parasite life cycle. The x-axis indicates the parasite stage and the y-axis (log_10_ scale) represents transcript fragments per kilobase of exon model per million reads mapped.

#### Expression of chimeric and mutant RACK1 proteins

To compare the ribosomal binding of human and *P. falciparum* RACK1, wild-type, chimeric, and mutant constructs were generated based on bioinformatics and structural model analyses (**FIGURE 2A**, **FIGURE 3, and** **FIGURE 5A**), as well as previous work^1,8–10,52^. While most of the ribosome facing loop residues appear conserved, previous work has shown that small single or double amino acid changes in key regions are able to significantly change binding^1,9,10,52^. Thus the RDK→DDE mutation was included as a control to show any potential reduction of binding (**FIGURE 2A** **MUTANT 3, and** **FIGURE 5A**). Previous work also emphasized the importance of the N-terminal region in RACK1 binding to the human 40S ribosomal subunit^1^. As such, we also generated chimeric RACK1 variants by exchanging the N-terminal regions of the HsRACK1 (residues 1-79) and PfRACK1 (residues 1-88). Multiple regions in the N-terminus of RACK1 differ between human and *P. falciparum* potentially influencing binding to ribosomes. Some alterations appear subtle, such as the introduction of a negatively charge aspartate, which may reduce interaction with the negatively charged rRNA backbone (**FIGURE 2A** **MUTANT 1**). The other differences appear more drastic in sequence of PfRACK1 protein, incorporating significant amounts of positively or negatively charged residue in contrast to the human homolog (**FIGURE 2A** **MUTANTS 2, 4, 5**). Therefore, these regions were exchanged between human and parasite RACK1 as well. It is important to note that the *P. falciparum* homolog is encoded using a significant number of rare human codons and therefore codon optimization is necessary for expression in mammalian cell lines. These constructs were expressed in HAP1 ΔRACK1 cells and *P. falciparum* parasites (**FIGURE 5**). RACK1 constructs with an N-terminal flag tag were transduced by lentivirus into HAP1 ΔRACK1 cells (**FIGURE 5B**). RACK1 was C-terminally tagged with fluorescent mNeonGreen followed by a 3x hemagglutinin affinity tag and electroporated into *P. falciparum* Dd2 cells (**FIGURE 5C**). Western blot analysis of human and parasite RACK1 lines indicates single band at ∼37 kDa for expression in human HAP1 cells and an ∼65 kDa band for expression in P. falciparum cells. The wild type proteins showed similar expression in both HAP1 and parasite cells (**FIGURE 5B-C**). Mutants and chimeric proteins varied in their expression between HAP1 (**FIGURE 5B-C, SUPPLEMENTAL FIGURE 2**). The patterns also differed between human and parasite cells. The constant between the two organisms was that the RDK to DDE mutant was anticipated to have significantly impaired binding in both organisms given the sequence conservation and its importance in previous work^1,9,10,52^. RACK1, like other ribosomal proteins^55^, is unstable when not bound to the ribosome^9^ and this, along with potential issued of folding, may account for the reduction in protein of some constructs.

**FIGURE 5.**
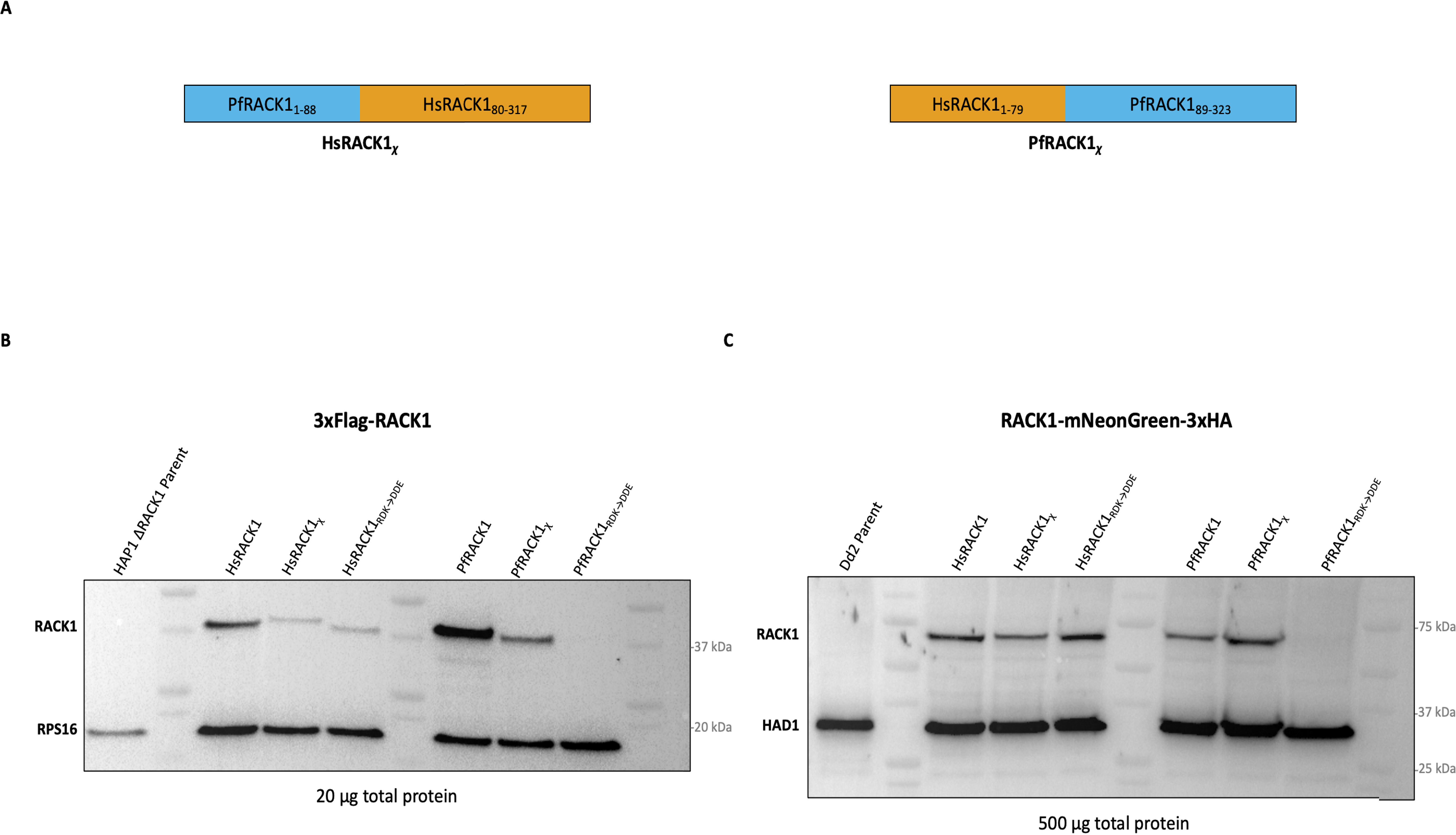
EXPRESSION OF RACK1 VARIANTS IN MAMMALIAN HAP1 ΔRACK1 CELLS AND *P. FALCIPARUM* PARASITES. **(A)** Schematic constructs for RACK1 variants. HsRACK1_χ_ designates the RACK1 chimera comprised of an N-terminus with PfRACK1 amino acids 1-88 followed by a c-terminus with HsRACK1 residues 80-317 (left). PfRACK1_χ_ designates the RACK1 chimera comprised of an N-terminus with HsRACK1 amino acids 1-79 followed by a c-terminus with PfRACK1 residues 89-323 (right). HsRACK1_RDK→DDE_ and PfRACK1_RDK→DDE_ constructs generated by mutating HsRACK1 Arg_36_/Lys_38_ or PfRACK1 Arg_43_/Lys_45_ to Asp/Glu **(B)** Western blot using anti-Flag antibody of HAP1 ΔRACK1 cell expressing RACK1 variants. 40S ribosomal protein 16 (uS9) blotted as control. **(C)** Western blot using anti-HA antibody of *P. falciparum* parasites expressing RACK1 variants. Haloacid dehalogenase-like hydrolase (HAD) blotted as control.

### Localization of RACK1 protein in *P. falciparum* cells

Previous work has stated that in *P. falciparum*, RACK1 is dispersed in the parasite cytoplasm during the schizont stage^20^ and it might be exported into the red blood cell^21^, the stage from which all previous ribosome structural data is based^22,23^. To reevaluate this phenomenon, live imaging microscopy was performed on *P. falciparum* Dd2 lines expressing C-terminally tagged mNeonGreen wild-type PfRACK. Parasites were treated with E64 to obtain late-stage schizonts. Hoechst 33342 stain (blue) was used to stain parasite DNA while BODIPY TR ceremide (red) was used to label host/parasite membrane, and mNeonGreen (green) was used to tag the exogenously expressed parasite RACK1 protein. Images were taken of the Dd2 parent line (**FIGURE 6A**) and Dd2 expressing mNeonGreen-tagged PfRACK1 (**FIGURE 6B**). BODIPY TR ceramide staining shows parasite daughter cells labeled within the host RBC while PfRACK1 mNeonGreen signal maintains close contact within these daughter cells showing no staining in the RBC cytoplasm. Therefore, imaging of schizonts did not show dispersal of RACK1 into the red blood cell cytoplasm, but rather the RACK1 protein remain in the parasite cytoplasm surrounding the daughter cells.

**FIGURE 6.**
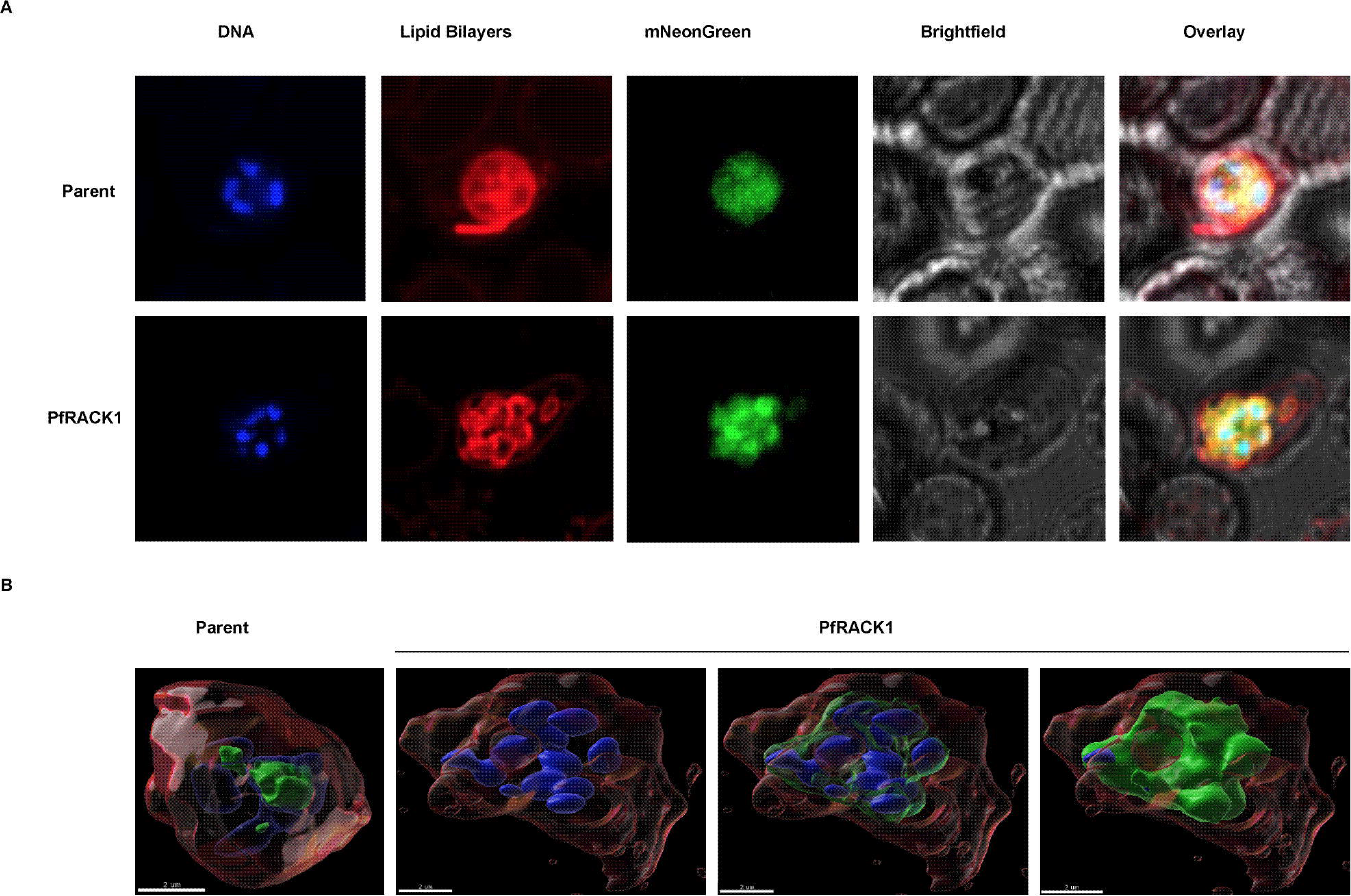
AIRYSCAN CONFOCAL FLUORESCENCE IMAGING MICROSCOPY OF MNEONGREEN-TAGGED PFRACK1 EXPRESSED IN *P. FALCIPARUM*. **(A)** Hoechst Stain (DNA), mNeonGreen (RACK1), BODIPY TR ceramide (lipid bilayers). Top) Dd2 parent line. Bottom) Dd2 parent line expressing mNeonGreen-tagged PfRACK1. **(B)** Two-dimensional overlay using Imaris software analysis of Airyscan images. Left) Dd2 parent line. Right) Dd2 parent line expressing mNeonGreen-tagged PfRACK1.

### Ribosome association of RACK1 chimeras and variants

To assess the binding of RACK1 to human and *P. falciparum* ribosomes and 40S subunits, we isolated crude ribosomes by running cell lysates through a sucrose cushion and then examined the presence of RACK1 in the crude ribosome pellet. The RACK1 variants expressed in HAP1 cells harboring the human RACK1 N-terminal region were able to bind the human ribosome while those that had the parasite RACK1 N-terminal region did not show significant binding to human ribosomes (**FIGURE 7A**). This difference in ribosome binding and potential association is most prominent between HsRACK1 and HsRACK1χ chimera as well as between PfRACK1 and PfRACK1χ chimera. HsRACK1 and PfRACK1χ chimera (human N-terminal region) were detected more in pelleted ribosome fraction than in the human cell lysate. PfRACK1 and HsRACK1χ chimera (*P.falciparum* N-terminal region) on the contrary, had opposite distribution, more protein was detected in lysates than in ribosomal pellets (**FIGURE 7A**). The loss of binding to human ribosomes was observed for the RDK→DDE mutation as in previous studies^1,8,11,52^ (**FIGURE 7A**). Interestingly, all variants appeared to bind in *P. falciparum* parasite, potentially even the poorly expressed PfRACK1 RDK→DDE mutant, suggesting a difference in RACK1::40S subunit binding in the parasite (FIGURE 7B). Furthermore, Western blot analysis of ribosomal fractions collected during polysome profiling in parasites suggests that PfRACK1 is bound to the ribosome during translation (**SUPPLEMENTAL FIGURE 3**). Finally, the ribosome binding of RACK1 variants (Figure 7B) was also seen in by RT-qPCR analyses of rRNAs bound to the immune-precipitated RACK1 protein variants (**SUPPLEMENTAL FIGURE 4**). Immunoprecipitation of all RACK1 variants and chimeras, except PfRACK1 RDK→DDE mutant, resulted in enrichment of bound rRNAs in RT-qPCR analysis (**SUPPLEMENTAL FIGURE 4**). As such, our data suggests that in comparison to the human RACK1 protein PfRACK1, binds specifically to the parasite ribosomes with little, if any, affinity towards human ribosomes.

**FIGURE 7.**
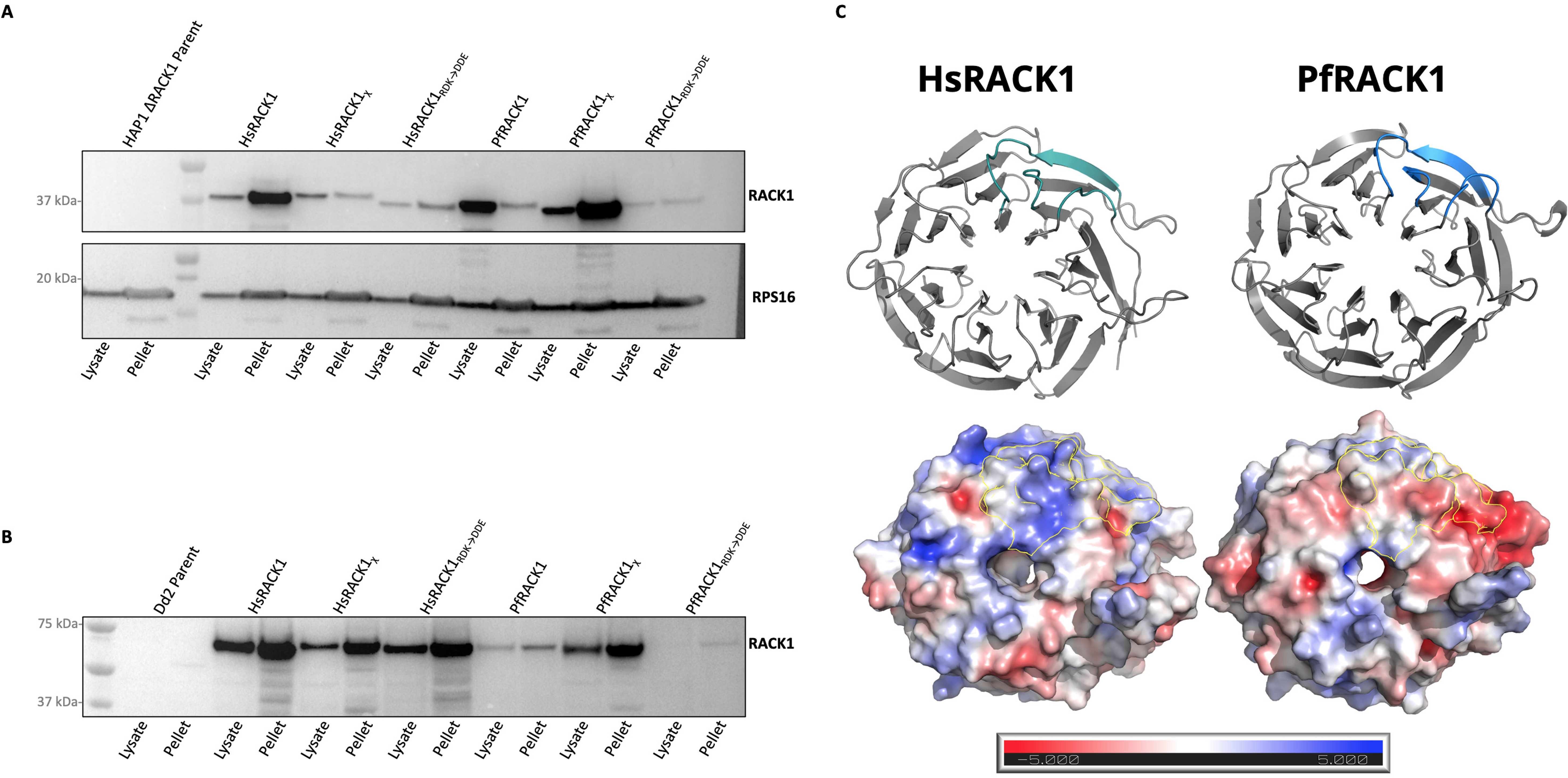
RIBOSOMAL BINDING OF RACK1 VARIANTS IN MAMMALIAN HAP1 ΔRACK1 CELLS AND *P. FALCIPARUM* PARASITES. **(A)** Western blot of lysates and post-sucrose cushion centrifugation pellets of HAP1 ΔRACK1 cells expressing N-terminally 3xFlag tagged RACK1 variants. 40S ribosomal protein 16 (uS9) blotted as control. **(B)** Western blot of lysates and post-sucrose cushion centrifugation pellets from *P. falciparum* parasites expressing C-terminally 3xHA-tagged RACK1 variants. **(C)** Ribbon model (above) displays ribosome facing structure with colored region indicating N-terminal section previously shown to be important for binding in mammalian cells. Electrostatic map (below) of above ribbon model ribosome-facing surface for human and *P. falciparum* RACK1 proteins generated using PyMOL APBS Electrostatics plugin. Outlines indicate colored region of N-terminal section previously shown to be important for RACK1 binding in mammalian cells.

## Discussion

With an approximately 60% shared sequence identity between the human and *P. falciparum* RACK1 proteins, sequence, and structural analysis reveal that PfRACK1 maintains all the hallmark characteristics of homologs where ribosomal binding is seen (**FIGURES 1–3**). This is particularly true of ribosomes facing residues. Combining the sequence analysis data with the *de novo* structure of PfRACK1 modeled onto the parasite 80S ribosome, most of the ribosome facing loop residues appear conserved and similar to human RACK1 protein (**FIGURES 2 and 3**). Additionally, the potential rRNA interactions also appear to be conserved in the parasite, as do those of RPS16 (uS9), RPS17 (eS17), and RPS3 (uS3) except for a potential alteration to the RPS3 C-terminal disordered region (**SUPPLEMENTAL FIGURE 1**). Bioinformatics analysis of PfRACK1^20,56^ mRNA abundance, demonstrates that the transcriptional profile of PfRACK1 mRNA follows other ribosomal proteins (**FIGURE 4**), further enabling its role in protein synthesis. Taken together, this data as well as previous studies on yeast and mammalian RACK1 homologs, suggested that PfRACK1 should be able to bind to the parasite 40S ribosomal subunit.

To examine PfRACK1 binding of the 40S ribosome subunit we generated RACK1 variants based on bioinformatics and structural analysis. We also generated chimeras between human and *P. falciparum* RACK1 proteins, switching N-terminal regions. Expression of the wild-type HsRACK1 and PfRACK1 appeared to be comparable within HAP1 ΔRACK1 mammalian cells and *P. falciparum* parasites (**FIGURE 5B-C**). In parasite lines, expression of all variants appeared approximately equal with the exception of the PfRACK1_RDK→DDE_ mutant that is expected to have reduced ribosome binding. In the mammalian line, both human and *P. falciparum* RACK1_RDK→DDE_ mutants, and RACK1 chimera with N-terminal region from *P. falciparum* (HsRACK1χ) showed reduced expression. It has been shown that, like most ribosomal proteins^55^, RACK1 becomes unstable when not bound to the ribosome^9^, which may explain the lower expression levels and predict subsequent ribosome binding in that organism. In this respect, those RACK1 variants that contained the *P. falciparum* N-terminal region and those with the RDK→DDE mutation might have reduced stability arguing for some differences and specificity in RACK1 binding in *P. falciparum* cells.

Since it was previously suggested that PfRACK1 might be exported during the schizont stage into the red blood cell cytoplasm^21^ we tagged PfRACK1 with the fluorescent reporter mNeonGreen. To enrich for parasites during the schizont stage, parasites were treated with E64, which prevents RBC membrane rupture^48,49^. Parasites were subsequently examined by confocal microscopy with airyscan to determine PfRACK1 localization in relation to parasite DNA and lipid bilayers. We did not find confirmation for the previously observed phenomenon of PfRACK1 export^21^, but rather confirmation of the localization and dispersion of PfRACK1 in the parasitic cytoplasm^20^. Our data indicated that mNeonGreen-tagged PfRACK1 remained within the schizont cytoplasm (**FIGURE 6**). While our data suggest that PfRACK1 is found within the parasite during the schizont stage, it is possible that the parasites were not incubated long enough to see release of the protein into the RBC cytoplasm. However, any phenomenon of exported PfRACK1 protein is most likely due to the perforation of the parasitophorous vacuole that occurs at late in this stage^57^.

Finally, our data indicates that P. falciparum RACK1 protein may have a different mode of binding to parasite ribosomes (**FIGURE 7C/8**). Previous studies reported that yeast RACK1 homolog Asc1 (ScAsc1) had been shown to be able to bind the human 40S ribosome subunit^8^. Although PfRACK1 shares a higher identity with HsRACK1 than ScAsc1 (59.55 % vs. 53.82%), it could not bind the human ribosome (**FIGURE 7A**). Analysis of ribosome binding in the parasite cells, however, shows that not only does PfRACK1 bind the ribosome (**FIGURE 7B**, **SUPPLEMENTAL FIGURE 3 and 4**), but that also binding is altered from that of human ribosomes (**FIGURE 7B**, S**UPPLEMENTAL FIGURE 3 and 4**), with less emphasis on N-terminus driven binding in parasite. Polysome profiling data also show that PfRACK1 is indeed found in the 80S and polysome fractions (**SUPPLEMENTAL FIGURES 3 and 4**). Still there is a reduction in the presence of PfRACK1 in this faction compared to the HsRACK1 and PfRACK1_χ_ variants when assayed in human cells. This again suggests an alteration in PfRACK1 binding of the 40S subunit and species-specific interactions^9–11,53,54^. Using the previously generated cryoEM structure of HsRACK1 and our *de novo* model of PfRACK1, we modeled the electrostatic composition of the ribosome-facing surface of RACK1 for human and *P. falciparum* (**FIGURE 7C**). The comparison shows a dramatic difference in the charge distribution between the two organisms, particularly in the N-terminal region thought to be most important for binding in humans (**FIGURE 7C, OUTLINED REGION**). This once again suggests a difference in mode of RACK1 binding in parasites versus the human host. However, this data does not include any post-translational modifications that could further alter ribosome binding of RACK1 in parasite.

According to currently available structural data, RACK1 does not copurify with schizont 80S ribosomes^22,23^. This may result from purification method artifacts^58^, such as the loss of polysomes with saponin lysis^59^. The presence of RACK1 on the ribosome seen in crude ribosomes and polysome profiling data here (**FIGURE 7B**, **SUPPLEMENTAL FIGURES 3 and 4)** and from others^60^ indicate that PfRACK1 does indeed bind the ribosome. However, this does not exclude the possibility of a change in PfRACK1::40S ribosome binding dynamics during the schizont stage. It is important to note that there is an almost log_10_ reduction in PfRACK1 transcript expression in the schizont stage (**FIGURE 4**). One could hypothesize that the drastic reduction of PfRACK1 expression and ribosome binding, which has previously been shown to globally downregulate mRNA translation^9^, would be energetically favorable as the parasite prepares to release short-lived merozoites.

## Acknowledgements

We thank Slavica Pavlovic Djuranovic, and members of Daniel Goldberg’s and Sergej Djuranovic’s lab for helpful comments. This work is supported by NIH R01 GM112824, NIH R01 GM136823 and NIMH R01 MH116999 to SD, NIH T32 GM007067 and NIMH R01 MH116999 to JE. We are also thankful to WUCCI center of Washington University on the help with this project. The Washington University LEAP Gap Fund supported this project through the Skandalaris Center for Interdisciplinary Innovation and Entrepreneurship under award number #1014, the Washington University Institute of Clinical and Translational Sciences, and the NIH/National Center for Advancing Translational Sciences (NCATS) grant UL1TR002345. The authors declare that they have no competing interests.

## References

1. Adams, D. R., Ron, D. & Kiely, P. A. RACK1, A multifaceted scaffolding protein: Structure and function. Cell Commun. Signal. 9, 22 (2011).

2. Jain, B. P. & Pandey, S. WD40 Repeat Proteins: Signalling Scaffold with Diverse Functions. Protein J. 37, 391–406 (2018).

3. Li, D. & Roberts, R. Human Genome and Diseases: WD-repeat proteins: structure characteristics, biological function, and their involvement in human diseases. Cell. Mol. Life Sci. 58, 2085–2097 (2001).

4. Ron, D. et al. Cloning of an intracellular receptor for protein kinase C: a homolog of the beta subunit of G proteins. Proc. Natl. Acad. Sci. 91, 839–843 (1994).

5. Yuan, L. et al. A RACK1-like protein regulates hyphal morphogenesis, root entry and in vivo virulence in Verticillium dahliae. Fungal Genet. Biol. 99, 52–61 (2017).

6. Bradford, W. et al. Eukaryotic G Protein Signaling Evolved to Require G Protein-Coupled Receptors for Activation. Sci. Signal. 6, ra37–ra37 (2013).

7. Smith, B. L. & Mochly-Rosen, D. Inhibition of protein kinase C function by injection of intracellular receptors for the enzyme. Biochem. Biophys. Res. Commun. 188, 1235–1240 (1992).

8. Johnson, A. G. et al. RACK1 on and off the ribosome. RNA 25, 881–895 (2019).

9. Gallo, S. et al. RACK1 Specifically Regulates Translation through Its Binding to Ribosomes. Mol. Cell. Biol. 38, 1–16 (2018).

10. Sengupta, J. et al. Identification of the versatile scaffold protein RACK1 on the eukaryotic ribosome by cryo-EM. Nat. Struct. Mol. Biol. 11, 957–962 (2004).

11. Thompson, M. K., Rojas-Duran, M. F., Gangaramani, P. & Gilbert, W. V. The ribosomal protein Asc1/RACK1 is required for efficient translation of short mRNAs. Elife 5, 1–20 (2016).

12. Nilsson, J., Sengupta, J., Frank, J. & Nissen, P. Regulation of eukaryotic translation by the RACK1 protein: a platform for signalling molecules on the ribosome. EMBO Rep. 5, 1137–1141 (2004).

13. Kouba, T., Rutkai, E., Karásková, M. & Valášek, L. S. The eIF3c/NIP1 PCI domain interacts with RNA and RACK1/ASC1 and promotes assembly of translation preinitiation complexes. Nucleic Acids Res. 40, 2683–2699 (2012).

14. Ceci, M. et al. Release of eIF6 (p27BBP) from the 60S subunit allows 80S ribosome assembly. Nature 426, 579–584 (2003).

15. Sugiyama, T. et al. Sequential Ubiquitination of Ribosomal Protein uS3 Triggers the Degradation of Non-functional 18S rRNA. Cell Rep. 26, 3400–3415.e7 (2019).

16. Joazeiro, C. A. P. Ribosomal Stalling During Translation: Providing Substrates for Ribosome-Associated Protein Quality Control. Annu. Rev. Cell Dev. Biol. 33, 343–368 (2017).

17. Kuroha, K. et al. Receptor for activated C kinase 1 stimulates nascent polypeptide-dependent translation arrest. EMBO Rep. 11, 956–961 (2010).

18. Ikeuchi, K., Yazaki, E., Kudo, K. & Inada, T. Conserved functions of human Pelota in mRNA quality control of nonstop mRNA. FEBS Lett. 590, 3254–3263 (2016).

19. Sundaramoorthy, E. et al. ZNF598 and RACK1 Regulate Mammalian Ribosome-Associated Quality Control Function by Mediating Regulatory 40S Ribosomal Ubiquitylation. Mol. Cell 65, 751–760.e4 (2017).

20. Blomqvist, K., DiPetrillo, C., Streva, V. A., Pine, S. & Dvorin, J. D. Receptor for Activated C-Kinase 1 (PfRACK1) is required for Plasmodium falciparum intra-erythrocytic proliferation. Mol. Biochem. Parasitol. 211, 62–66 (2017).

21. Vincensini, L. et al. Proteomic Analysis Identifies Novel Proteins of the Maurer’s Clefts, a Secretory Compartment Delivering Plasmodium falciparum Proteins to the Surface of Its Host Cell. Mol. Cell. Proteomics 4, 582–593 (2005).

22. Wong, W. et al. Cryo-EM structure of the Plasmodium falciparum 80S ribosome bound to the anti-protozoan drug emetine. Elife 3, (2014).

23. Sun, M. et al. Dynamical features of the Plasmodium falciparum ribosome during translation. Nucleic Acids Res. 43, gkv991 (2015).

24. Bunnik, E. M. et al. The mRNA-bound proteome of the human malaria parasite Plasmodium falciparum. Genome Biol. 17, 1–18 (2016).

25. Aurrecoechea, C. et al. PlasmoDB: a functional genomic database for malaria parasites. Nucleic Acids Res. 37, D539–D543 (2009).

26. López-Barragán, M. J. et al. Directional gene expression and antisense transcripts in sexual and asexual stages of Plasmodium falciparum. BMC Genomics 12, 587 (2011).

27. Zanghì, G. et al. A Specific PfEMP1 Is Expressed in P. falciparum Sporozoites and Plays a Role in Hepatocyte Infection. Cell Rep. 22, 2951–2963 (2018).

28. Ruiz Carrillo, D. et al. Structure of human Rack1 protein at a resolution of 2.45 Å. Acta Crystallogr. Sect. F Struct. Biol. Cryst. Commun. 68, 867–872 (2012).

29. Altschul, S. F. et al. Protein database searches using compositionally adjusted substitution matrices. FEBS J. 272, 5101–5109 (2005).

30. Altschul, S. Gapped BLAST and PSI-BLAST: a new generation of protein database search programs. Nucleic Acids Res. 25, 3389–3402 (1997).

31. Zimmermann, L. et al. A Completely Reimplemented MPI Bioinformatics Toolkit with a New HHpred Server at its Core. J. Mol. Biol. 430, 2237–2243 (2018).

32. Klausen, M. S. et al. NetSurfP-2.0: Improved prediction of protein structural features by integrated deep learning. Proteins Struct. Funct. Bioinforma. 87, 520–527 (2019).

33. Hanson, J., Yang, Y., Paliwal, K. & Zhou, Y. Improving protein disorder prediction by deep bidirectional long short-term memory recurrent neural networks. Bioinformatics btw678 (2016). doi:10.1093/bioinformatics/btw678

34. Jones, D. T. & Cozzetto, D. DISOPRED3: precise disordered region predictions with annotated protein-binding activity. Bioinformatics 31, 857–863 (2015).

35. Yan, R., Xu, D., Yang, J., Walker, S. & Zhang, Y. A comparative assessment and analysis of 20 representative sequence alignment methods for protein structure prediction. Sci. Rep. 3, 2619 (2013).

36. Heffernan, R., Yang, Y., Paliwal, K. & Zhou, Y. Capturing non-local interactions by long short-term memory bidirectional recurrent neural networks for improving prediction of protein secondary structure, backbone angles, contact numbers and solvent accessibility. Bioinformatics 33, 2842–2849 (2017).

37. Jones, D. T. Protein secondary structure prediction based on position-specific scoring matrices 1 1Edited by G. Von Heijne. J. Mol. Biol. 292, 195–202 (1999).

38. Schrodinger, L. The PyMOL Molecular Graphics Development Component, Version 2.4. (2020).

39. Brown, A., Shao, S., Murray, J., Hegde, R. S. & Ramakrishnan, V. Structural basis for stop codon recognition in eukaryotes. Nature 524, 493–496 (2015).

40. Jurrus, E. et al. Improvements to the APBS biomolecular solvation software suite. Protein Sci. 27, 112–128 (2018).

41. Waterhouse, A. et al. SWISS-MODEL: homology modelling of protein structures and complexes. Nucleic Acids Res. 46, W296–W303 (2018).

42. Bienert, S. et al. The SWISS-MODEL Repository—new features and functionality. Nucleic Acids Res. 45, D313–D319 (2017).

43. Guex, N., Peitsch, M. C. & Schwede, T. Automated comparative protein structure modeling with SWISS-MODEL and Swiss-PdbViewer: A historical perspective. Electrophoresis 30, S162–S173 (2009).

44. Studer, G. et al. QMEANDisCo—distance constraints applied on model quality estimation. Bioinformatics 36, 1765–1771 (2020).

45. Bertoni, M., Kiefer, F., Biasini, M., Bordoli, L. & Schwede, T. Modeling protein quaternary structure of homo- and hetero-oligomers beyond binary interactions by homology. Sci. Rep. 7, 10480 (2017).

46. Trager, W. & Jensen, J. B. Human malaria parasites in continuous culture. 1976. J. Parasitol. 91, 484–6 (2005).

47. Fidock, D. A. & Wellems, T. E. Transformation with human dihydrofolate reductase renders malaria parasites insensitive to WR99210 but does not affect the intrinsic activity of proguanil. Proc. Natl. Acad. Sci. 94, 10931–10936 (1997).

48. Salmon, B. L. From the Cover: Malaria parasite exit from the host erythrocyte: A two-step process requiring extraerythrocytic proteolysis. Proc. Natl. Acad. Sci. 98, 271–276 (2001).

49. Glushakova, S., Mazar, J., Hohmann-Marriott, M. F., Hama, E. & Zimmerberg, J. Irreversible effect of cysteine protease inhibitors on the release of malaria parasites from infected erythrocytes. Cell. Microbiol. 11, 95–105 (2009).

50. McGlincy, N. J. & Ingolia, N. T. Transcriptome-wide measurement of translation by ribosome profiling. Methods 126, 112–129 (2017).

51. Arthur, L. L. et al. Translational control by lysine-encoding A-rich sequences. Sci. Adv. 1, e1500154 (2015).

52. Coyle, S. M., Gilbert, W. V. & Doudna, J. A. Direct Link between RACK1 Function and Localization at the Ribosome In Vivo. Mol. Cell. Biol. 29, 1626–1634 (2009).

53. Rollins, M. G., Jha, S., Bartom, E. T. & Walsh, D. RACK1 evolved species-specific multifunctionality in translational control through sequence plasticity within a loop domain. J. Cell Sci. 132, jcs228908 (2019).

54. Nielsen, M. H., Flygaard, R. K. & Jenner, L. B. Structural analysis of ribosomal RACK1 and its role in translational control. Cell. Signal. 35, 272–281 (2017).

55. Warner, J. R. In the absence of ribosomal RNA synthesis, the ribosomal proteins of HeLa Cells are synthesized normally and degraded rapidly. J. Mol. Biol. 115, 315–333 (1977).

56. von Bohl, A. et al. A WD40-repeat protein unique to malaria parasites associates with adhesion protein complexes and is crucial for blood stage progeny. Malar. J. 14, 435 (2015).

57. Hale, V. L. et al. Parasitophorous vacuole poration precedes its rupture and rapid host erythrocyte cytoskeleton collapse in Plasmodium falciparum egress. Proc. Natl. Acad. Sci. 114, 3439–3444 (2017).

58. Regmi, S., Rothberg, K. G., Hubbard, J. G. & Ruben, L. The RACK1 signal anchor protein from Trypanosoma brucei associates with eukaryotic elongation factor 1A: A role for translational control in cytokinesis. Mol. Microbiol. 70, 724–745 (2008).

59. Lacsina, J. R., LaMonte, G., Nicchitta, C. V & Chi, J.-T. Polysome profiling of the malaria parasite Plasmodium falciparum. Mol. Biochem. Parasitol. 179, 42–46 (2011).

60. Caro, F., Ahyong, V., Betegon, M. & DeRisi, J. L. Genome-wide regulatory dynamics of translation in the Plasmodium falciparum asexual blood stages. Elife 3, 1–24 (2014).

