## Supplementary material for "Association of RACK1 protein with ribosomes in *Plasmodium falciparum*": Suplemental Figures 1-4

### RPS3 (uS3) Alignment

```
HsRPS3      MAVQISKKRKFVADGIFKAELNEFLTRELAEDGYSGVEVRVTPTRTEIIILATRTQNVLG     60
PfRPS3      MSAPISKKRKFINDGVFQAELNEFLARILAEDGYSGVEVRVTPIRTEVIIRATRTREVLG     60
            *:. *******: **:*:*******:* *************** ***:** ****::***

HsRPS3      EKGRRIRELTAVVQKRF-GFPEGSVELYAEKVATRGLCAIAQAESLRYKLLGGLAVRRAC     119
PfRPS3      DKGRRIRELTSLVQKRFFNKSTNSVELFAERVEHRGLCAMAQAESLRYKLLKGLAVRRAC     120
            :*********::***** .   .****:**:*  *****:*********** ********

HsRPS3      YGVLRFIMESGAKGCEVVVSGKLRGQRAKSMKFVDGLMIHSGDPVNYYVDTAVRHVLLRQ     179
PfRPS3      YGVLRHIMESGAKGCEVIVSGKLRAQRAKSMKFRDGYLISTGEPSKRFVNTATRSAQLKQ     180
            *****.***********:******.******** ** :* :*:* : :*:**.* . *:*

HsRPS3      GVLGIKVKIMLPWDPTGKIGPKKPLPDHVSIVEPKDEILPTTPISEQKGGKPEPPAMPQP     239
PfRPS3      GVLGIKVKIMLPTAIDTRTGLTSILPDNISVLEPKTDTVDL-------------------     221
            ************     : *  .. ***::*::*** : :

HsRPS3      VPTA         243
PfRPS3      ----         221
```

### RPS16 (uS9) Alignment

```
HsRPS16     MPSKGPLQSVQVFGRKKTATAVAHCKRGNGLIKVNGRPLEMIEPRTLQYKLLEPVLLLGK     60
PfRPS16     --MTTKVKRVQTFGKKKTAVAVATVTNGKGLIKLNGKNLDLVEPYILKTKVYEPLWLIGS     58
               .  :: **.**:****.***  ..*:****:**: *:::**  *: *: **: *:*.

HsRPS16     ERFAGVDIRVRVKGGGHVAQIYAIRQSISKALVAYYQKYVDEASKKEIKDILIQYDRTLL     120
PfRPS16     GKLKNLDIRIRVKGGGQTSQIYAIRQAIGKGIISYYQKYVDESTKKELKDVLLRYDRSLL     118
             :: .:***:******:.:*******:.*.:::**********:***:**:*::***:**

HsRPS16     VADPRRCESKKFGGPGARARYQKSYR 146
PfRPS16     VGDTRRCEPKKFGGKGARARYQKSYR 144
            *.* **** ***** ***********
```

### RPS17 (eS17) Alignment

```
HsRPS17     MGRVRTKTVKKAARVIIEKYYTRLGNDFHTNKRVCEEIAIIPSKKLRNKIAGYVTHLMKR     60
PfRPS17     MGRVRTKTIKRAARQIVEKYYAKLTLDFQINKKITEEVAIIPSKRMKNKVAGFVTHLMKR     60
            ********:*:*** *:****::*  **: **::  **:******::.**:**:*******

HsRPS17     IQRGPVRGISIKLQEEERERRDNYVPEVSALDQEIIEVDPDTKEMLKLLDFGSLSNLQVT     120
PfRPS17     IQKGPVRGISLKLQEEERERRLDFVPEKSQIDVSVIYVEPDTLRMIKSLGIN-ISNMKVH     119
            **:*******:********** ::*** * :* .:* *:*** .*:* *.:. :**::*

HsRPS17     QPTVGMNFKTPRGPV--- 135
PfRPS17     NPMINTNQQKQNRMNNQF 137
            :* :. * :. .
```

**Supplemental Figure 1. Multiple sequence alignment of *H. sapiens* and *P. falciparum* RPS3 (uS3), RPS16 (uS9), RPS17 (eS17) proteins.** The *H. sapiens* and *P. falciparum* sequences were collected from the UniProt database[52] and PlasmoDB[31], respectively, and aligned using the Clustal Ω MSA tool[53]. Residues in the 3-5 Å range were determined using PyMol.

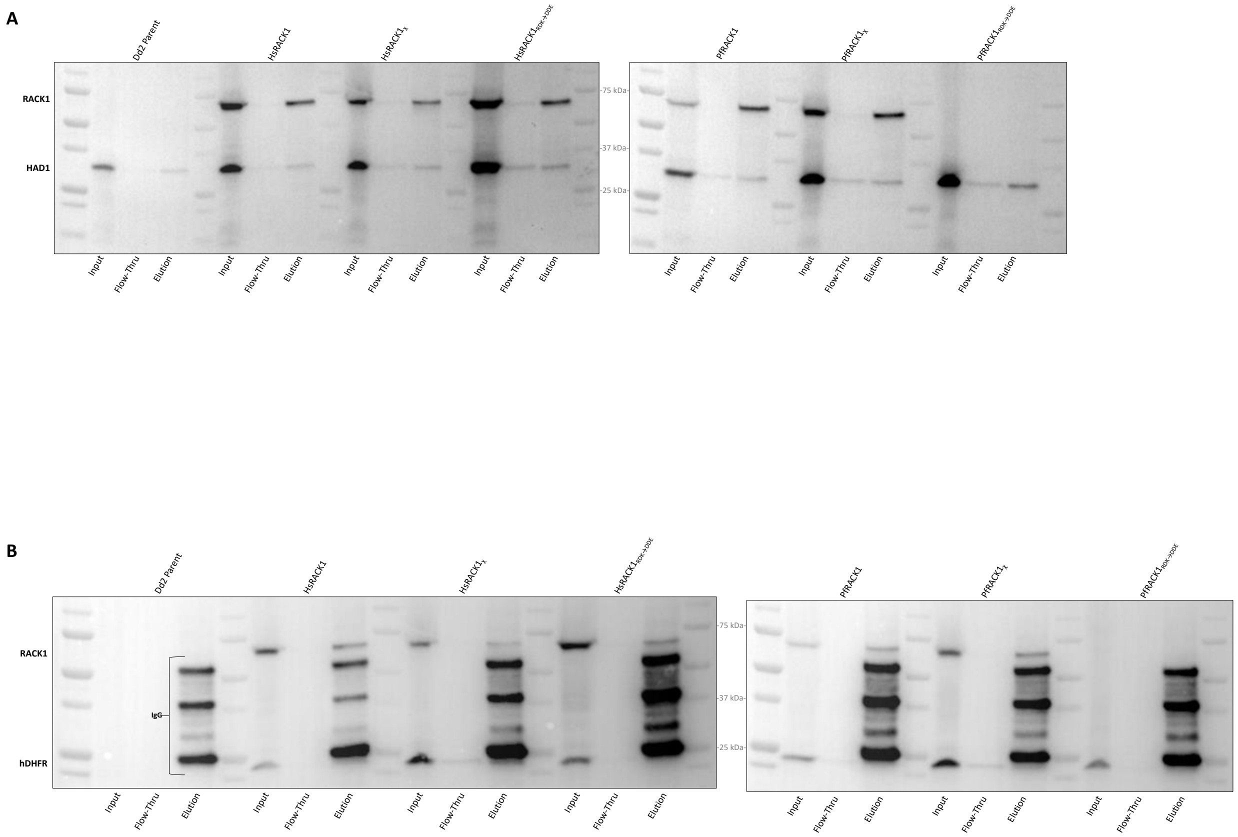


**SUPPLEMENTAL FIGURE 2. HA-IMMUNOPRECIPITATION OF RACK1 VARIANTS EXPRESSED IN *P. FALCIPARUM* DD2. (A)** Samples were blotted with mouse anti-HA-HRP antibody and rabbit anti-PfHAD1/anti-rabbit-HRP. **(B)** Samples were blotted with mouse anti-HA and mouse anti-hDHFR antibodies and subsequently blotted with anti-mouse-HRP antibody demonstrating presence of plasmid resistance containing PfRACK1$_{RDK \rightarrow DDE}$ mutant.

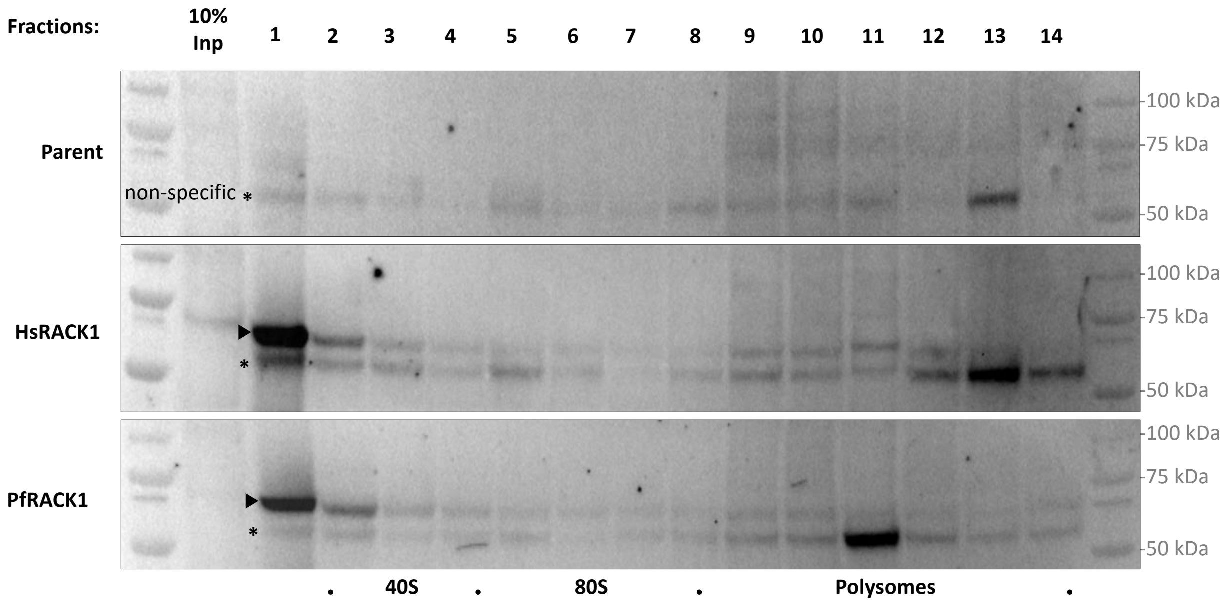


**Supplemental Figure 3. Polysome profiling of HsRACK1, PfRACK1, and PfRACK1$_\chi$ variants in *P. falciparum* Dd2 parasites.** Polysome profiling fractions generated from *P. falciparum* parasites expressing HsRACK1, PfRACK1, and PfRACK1$_\chi$ variants were analyzed. Western blot shows RACK1 variant polysome fraction localization. RT-qPCR analysis shows localization of 18S and 28S rRNAs associated with the 40S and 80S subunits, respectively. Arrowhead indicates non-specific band found in parent line not correlating to RACK1 variants.

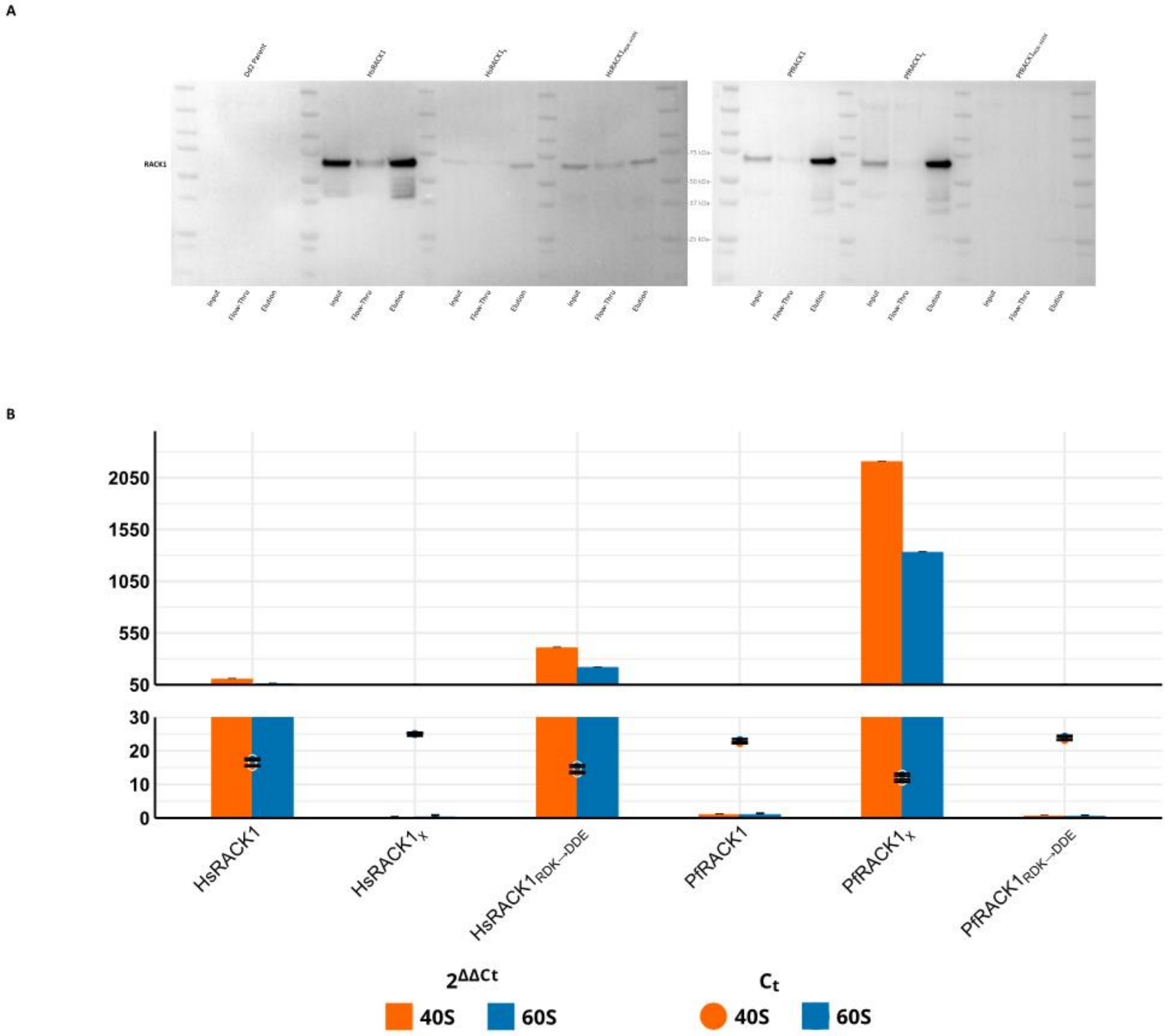


**Supplemental Figure 4. HA-immunoprecipitation of RACK1 variants bound to rRNA expressed in *P. falciparum* Dd2. (A)** Samples were blotted with mouse anti-HA-HRP antibody to show enrichment of HA-tagged RACK1 variants. **(B)** RT-qPCR was performed on RNA isolated from HA-immunoprecipitations of parasites expressing the RACK1 variants to examine rRNA binding. Samples were normalized to Dd2 parent line and then to the PfRACK1 wild type variant. The Ct (top) is displayed for each variant and the $2^{-\Delta\Delta Ct}$ (bottom) was calculated and plotted for each RACK1 variant.
